# Leveraging Supramolecular Polymers to Induce the Targeted Protein Degradation of α-Synuclein

**DOI:** 10.64898/2026.07.20.739556

**Authors:** Windfield S. Swetman, Malay Mondal, Ashe M. Davis, Vijay Rangachari, Tristan D. Clemons

**Affiliations:** School of Polymer Science and Engineering, University of Southern Mississippi, Hattiesburg, MS, 39406, USA; Department of Chemistry and Biochemistry, School of Mathematics and Natural Sciences, University of Southern Mississippi, Hattiesburg, MS, USA

**Keywords:** targeted protein degradation, neurodegenerative disease, peptide amphiphile, α-Synuclein, supramolecular polymers, Parkinson’s Disease

## Abstract

Halting the progression of neurodegenerative diseases remains one of the foremost challenges in medicinal chemistry due to the complex biology that drives disease progression. For example, a hallmark of synucleinopathies, such as Parkinson’s disease, is the misfolding and aggregation of the protein α-Synuclein (α-Syn), driving the formation of toxic oligomers and fibrils that avoid natural intracellular clearance mechanisms, participate in unusual protein-protein interactions, and ultimately contribute to the death of dopaminergic neurons. The field of targeted protein degradation (TPD) has emerged as an innovative therapeutic route to selectively degrade proteins of interest that leverage natural intracellular protein degradation machinery. First generation TPD therapeutics have traditionally been designed as bifunctional, chimeric compounds in which a short covalent linker tethers a ligand designed to bind target proteins to a ligand that initiates an either proteosome- or lysosome-dependent protein degradation cascade. While initial studies have indicated the promise of these approaches, translation to the clinical setting has been challenging due to difficulties in achieving cellular internalization, long-term stability, and establishment of a generalizable strategy. To overcome these obstacles, this work has focused on adding modularity and dynamic capability to this classical model by leveraging a multivalent macromolecular approach to TPD. Specifically, peptide amphiphiles (PAs) were designed to self-assemble into high-aspect-ratio supramolecular nanofibers and present peptide epitopes on the surface of the fibers to target simultaneous binding of α-Syn and recruitment of enzymes that facilitate entry into the lysosome-dependent chaperone-mediated autophagy protein degradation pathway. *In vitro* application of these bioactive PA nanofibers has demonstrated the ability to independently internalize in cells and reduce α-Syn protein levels selectively and effectively. While further optimization of this model has the potential to be a viable therapeutic against α-Syn aggregation, the modularity of these supramolecular nanofibers through facile monomer design and incorporation illustrates the potential of establishing a platform technology for targeting a diverse range of pathologic proteins.

**Graphical Abstract:** 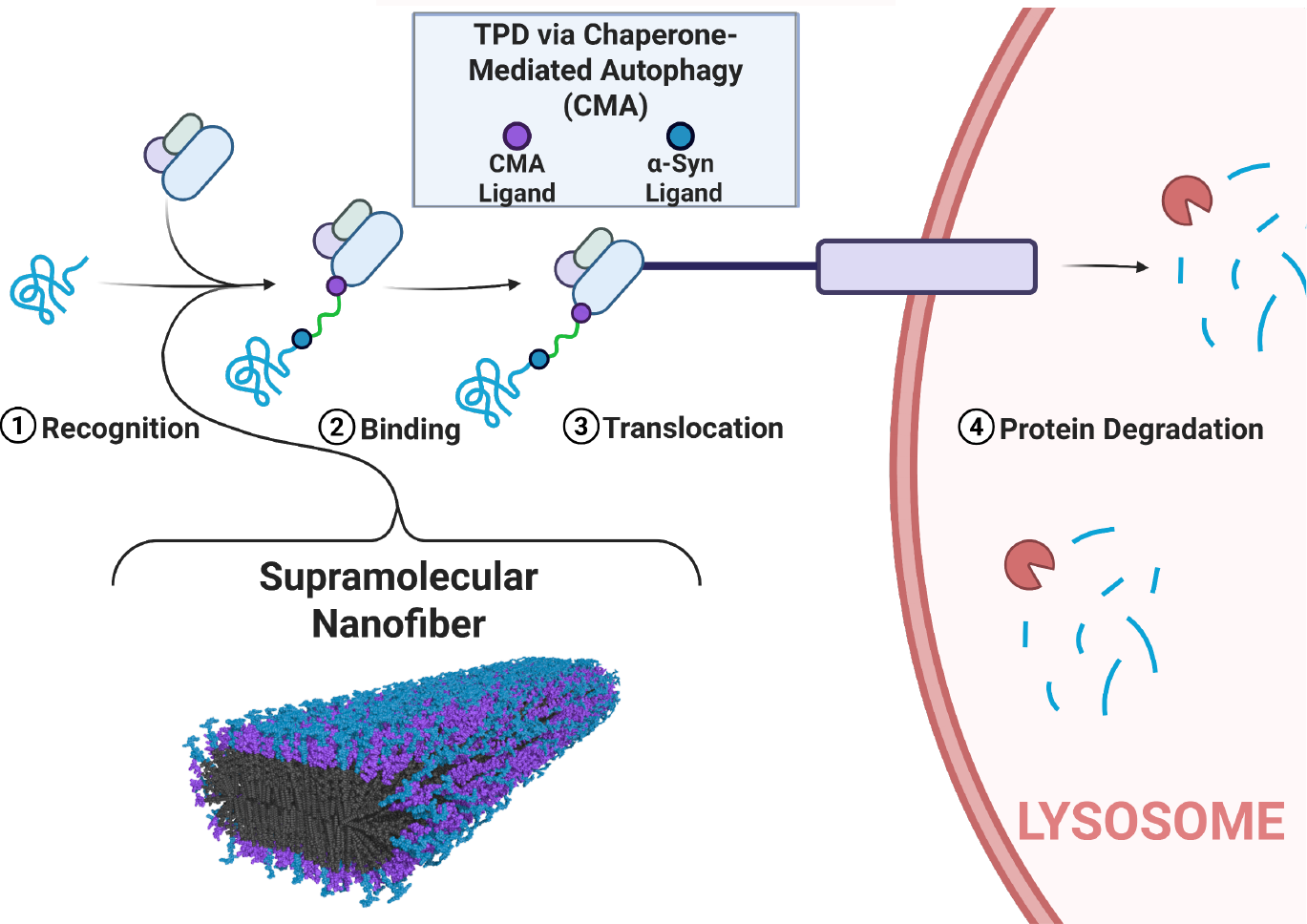

## Introduction

Neurodegenerative diseases encompass a class of disorders defined by the progressive loss of neuronal structure and function, ultimately leading to deteriorations in motor coordination, cognition, and quality of life. These diseases, including Alzheimer’s disease, Parkinson’s disease (PD), Huntington’s disease, and amyotrophic lateral sclerosis (ALS), often share pathological features such as the accumulation of misfolded proteins into toxic oligomeric and fibrillar aggregates.^1^ These toxic species participate in unusual protein-protein interactions and disrupt cellular homeostasis, triggering oxidative stress, mitochondrial dysfunction, neuroinflammation, and collectively resulting in neuronal death.^2-4^ Despite their growing societal impact, neurodegenerative diseases currently lack curative therapies, and existing treatments are primarily palliative.^3, 5^

A major obstacle in neurodegenerative disease treatment is that many pathogenic proteins are considered “undruggable” by conventional small-molecule approaches.^6^ These proteins lack deep hydrophobic pockets or catalytic domains typically exploited by inhibitors, rendering them resistant to modulation through classical binding-based pharmacology.^5^ In fact, fewer than 2% of disease-associated proteins are currently deemed druggable, highlighting the need for alternative modalities.^7^ One such protein is α-synuclein (α-Syn), an intrinsically disordered protein enriched in presynaptic terminals. Although widely recognized as a hallmark of PD, α-Syn has also been implicated in multiple other neurodegenerative diseases, including dementia with Lewy bodies, multiple system atrophy, and Alzheimer’s disease, where its misfolded and aggregated forms contribute to disease pathology.^3, 8-10^ Efforts to suppress α-Syn expression, block its aggregation, or promote its clearance have led to a range of strategies, including RNA interference, molecular chaperone modulation, and target protein regulation.^6,10-12^

Targeted protein degradation (TPD) offers a promising therapeutic strategy by utilizing endogenous cellular degradation systems to remove disease-driving proteins. Rather than inhibiting protein function, TPD can achieve complete elimination of the protein of interest (POI), providing a potentially more robust and long-lasting therapeutic effect. This approach bypasses the structural limitations of classically undruggable proteins and generally operates by leveraging either the ubiquitin-proteasome system (UPS) or lysosome-mediated pathways.^2, 5, 13, 14^ While the UPS is effective for soluble proteins of moderate size, lysosomal degradation pathways offer distinct advantages for eliminating aggregated, membrane-bound, or high-molecular-weight proteins that resist proteasomal processing.^15-17^

Among lysosomal mechanisms, chaperone-mediated autophagy (CMA) presents a particularly selective and energetically favorable route for achieving TPD. CMA operates through recognition of amino acid sequences, usually including the highly conserved pentapeptide KFERQ motif, within substrate proteins by the cytosolic chaperone Hsc70. The chaperone-substrate complex is trafficked to the lysosomal membrane, where it binds to the multimeric membrane protein LAMP-2A, enabling direct translocation of the substrate into the lysosomal lumen for degradation.^2, 15, 18-22^ Recent studies have demonstrated the feasibility of redirecting POIs to the CMA pathway via engineered KFERQ motif-bearing conjugates, enabling CMA-based TPD as a viable therapeutic platform.^15, 18, 22-24^

The vast majority of TPD strategies explored to date have leveraged a bifunctional, chimeric design linking a POI-targeting motif to an intracellular degradation signaling sequence.^2, 8, 25^ These chimeric degrader systems, regardless of mechanism, classically suffer from poor cellular uptake due to the increased molecular weight of these molecules and diminished efficacy against protein aggregates.^16^ Small molecule ligands have been investigated, but their use raises concerns regarding potential off-target effects.^16, 26, 27^ Peptide-based approaches offer enhanced selectivity and the potential for improved internalization, particularly when coupled with cell-penetrating peptides such as the trans-activator of transcription (TAT) peptide derived from the human immunodeficiency virus (HIV).^12, 18, 28^ However, these benefits are offset by rapid intracellular degradation and suboptimal pharmacokinetic profiles. Thus, there is a clear need for biocompatible, highly selective therapeutics that maintain proteolytic resistance and effectiveness in therapeutic applications.

Peptide amphiphiles (PAs) are peptide-based monomers capable of rapid self-assembly into elongated supramolecular nanostructures in aqueous environments. These assemblies are highly modular and can be designed to include a broad range of bioactive moieties, making them attractive for a broad range of biomedical applications. PAs that form high-aspect-ratio nanofibers at physiological pH typically consist of a hydrophobic alkyl tail conjugated to a peptide sequence with two domains: a beta-sheet–forming region to drive intermolecular cohesion between monomers, and a charged segment to ensure aqueous solubility. Self-assembly is initiated by the hydrophobic collapse of the alkyl tails into a nanofiber core, while beta-sheet formation stabilizes the nanostructure through intermolecular hydrogen bonding.^29-32^ Finally, a bioactive peptide sequence can be incorporated beyond the charged domain, to allow for peptide sequence presentation on the nanofiber surface. This modular PA design enables multivalent presentation of bioactive epitopes and the ability to co-assemble PA monomers within the same supramolecular polymer nanostructure to achieve multifunctional spatial and chemical bioactive display.^33-37^

The modular architecture of PA nanofibers enables control over assembly, stability, and functionality by varying the amino acid sequence, charge distribution, and hydrophobic regions. Recent studies have begun to explore covalent polymers and supramolecular nanostructures in the context of TPD, demonstrating that macromolecules such as nanospheres and liposomes can successfully induce degradation of intracellular proteins through proteasome- and lysosome-mediated pathways.^15, 23, 38-41^ Although PA nanofibers have yet to be applied for TPD, their inherent multivalency, stability, and biocompatibility make them a promising material platform for the development of next-generation degraders, particularly for directing aggregation-prone proteins toward lysosomal clearance pathways such as CMA.

## Discussion

### Peptide Amphiphile Nanofiber Characterization

To combine the benefits of dynamic, macromolecular architecture with the precision and target-specificity of TPD, we initially designed PA monomers that were capable of supramolecular co-assembly but individually target binding to α-Syn and trafficking into the CMA pathway, respectively. Efficient supramolecular co-assembly of functionalized PA monomers can be accomplished when PA monomers are built on the same backbone sequence and co-assembled into a randomly distributed copolymer.^32, 36, 42^ Both bioactive PA monomers used in this study were synthesized via conjugation of a palmitic acid to the N-terminus of the peptide sequence, VVAAEE. Using a diglycine linker to provide spacing between oppositely charged amino acids, the CMA-targeting PA was designed to present the highly conserved amino acid sequence, KFERQ, on the C-terminus and ultimately the surface of the co-assembled PA supramolecular polymers (CMA PA, Figure 1A). The peptide sequence, GVLYVGSKTR, derived from β-Synuclein, has been demonstrated to be an efficient binder of pathological α-Syn.^10, 12, 18, 28^ This sequence was chosen to be conjugated to the C-terminus of the α-Syn-targeting PA monomer to achieve α-Syn binding with our bioactive supramolecular polymers (α-Syn PA, Figure 1B). Due to the added hydrophobicity of this bioactive sequence, an additional two glutamic acid residues were added to this PA monomer design to ensure aqueous solubility was maintained to achieve self-assembly consistent with previous work on similar molecules.^43^ While neither was designed to induce the TPD of α-Syn as a homopolymer, co-assembly of these PA monomers into the same supramolecular polymer, produces a multivalent material capable of inducing both binding of α-Syn and recruitment of Hsc70 to induce effective lysosomal trafficking and, ultimately, targeted α-Syn degradation (Figure 1C).

**Figure 1.**
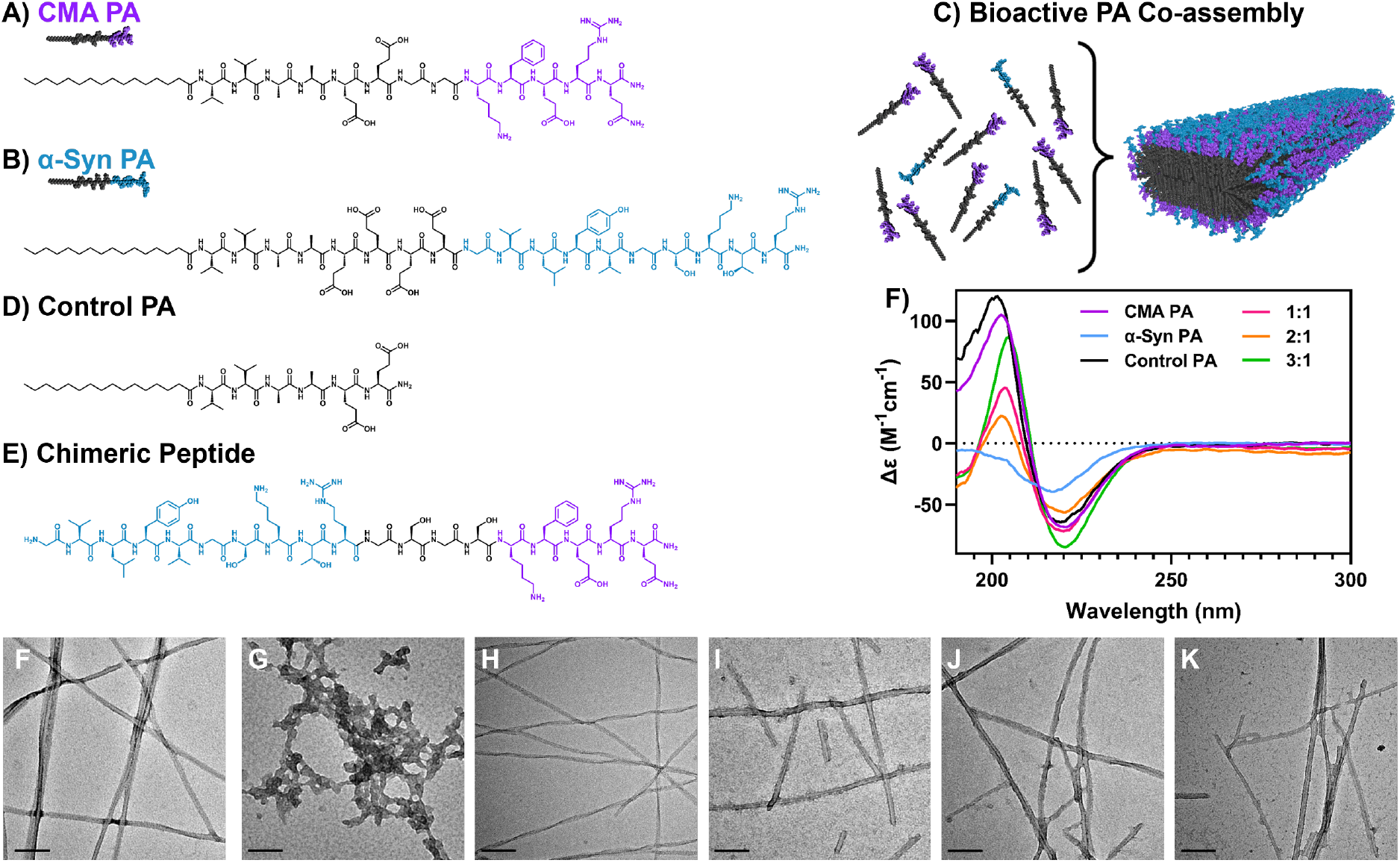
Design and Characterization of TPD Materials. A) CMA recruiting PA (CMA PA, purple). B) α-Syn binding PA (α-Syn PA, blue). C) Schematic representation of the 3:1 CMA: α-Syn bioactive PA supramolecular coassembly. D) control, fiber-forming PA without recruitment epitopes (Control PA). E) a bifunctional, chimeric peptide control (Chimeric Peptide). F) CD spectra displaying the secondary structure of the PA supramolecular polymers, including the homopolymers and co-assemblies between two bioactive PAs (CMA: α-Syn, mol%). TEM images of G) CMA PA homopolymer, H) α-Syn PA homopolymer, I) Control PA homopolymer, and three co-assembled supramolecular polymers of the bioactive PAs J) 1:1, K) 2:1, and L) 3:1 (CMA PA: α-Syn PA, mol%). All scale bars are 100 nm.

To study the effects of our TPD supramolecular polymers, we also synthesized two controls to assess the impact of macromolecular architecture and the importance of the bioactive ligands. A control PA nanofiber, without the inclusion of the bioactive epitopes for TPD was synthesized (Control PA, Figure 1D), and a chimeric peptide combining the two respective bioactive sequences presented from the bioactive PAs into a single peptide sequence, tethered with a glycine-serine (GSGS) spacer region, to function as a traditional chimeric TPD compound (Chimeric Peptide, Figure 1E). All PAs and the chimeric peptide were synthesized via solid-phase peptide synthesis, and the purity and identity of final, purified peptides were confirmed utilizing electrospray-ionization mass spectrometry (See supporting information for complete methods and Figure S1 for mass spectrometry purity assessment).

Initially, a combination of characterization techniques was used to probe the formation and morphology of the PA nanofibers. The critical aggregation concentration (CAC) of the PA monomers as both homo- and co-assemblies was assessed with a Nile red aggregation assay. The Nile red assay is based on the solvatochromic properties of Nile red, a lipophilic dye, which has weak flourescence in aqueous environments but intense fluorescence in hydrophobic environments, such as the core of the assembled nanofibers.^44^ The CAC for all formulations indicated a propensity for the PA monomers to form supramolecular assemblies in aqueous environments with CACs ranging from 0.7 to 6.4 μM (Figure S2). The formation of ordered, elongated PA nanofibers is dependent on strong intermolecular cohesion between monomers, where hydrophobic amino acid residues (i.e. valine and alanine residues) in our system are included for strong beta-sheet hydrogen bonding between monomers. To assess this secondary structure of the PA nanofibers, circular dichroism (CD) was used to confirm the presence of beta-sheet hydrogen bonding (Figure 1F). CD confirms the presence of beta-sheet hydrogen bonding in all supramolecular polymer assemblies, with the characteristic minima of 195 nm and maxima of 218 nm for beta-sheet hydrogen bonding observed.^34, 36, 45^. The homopolymer of the α-Syn PA, however, demonstrated a weak beta-sheet signature, likely a result of the additional hydrophobicity of this bioactive epitope, disrupting nanofiber formation. Transmission electron microscopy (TEM) enabled morphological assessment at the nanoscale of the assembled PA nanofibers (Figure 1G-L). Importantly, as a control, no ordered self-assembly was observed for the chimeric peptide as evidenced by Nile red assay, CD, and TEM (Figure S3). While the relatively long and hydrophobic bioactive epitope incorporated into the α-Syn PA design appears to disrupt nanofiber formation as a homopolymer, homopolymers of the CMA-targeting PA and the control PA self-assemble into high-aspect-ratio nanofibers with beta-sheet character, as evidenced by TEM and CD respectively. Importantly, co-assembly of the two bioactive PAs consisting of at least 50 mol% CMA PA also exhibit nanofiber formation with strong beta-sheet hydrogen bonding. This co-assembly strategy was utilized to create bifunctional, multivalent, supramolecular polymer nanofibers with the facile capability of controlling bioactive ligand ratios within the nanofiber to ultimately assess the importance of this characteristic on bioactive supramolecular polymer TPD efficacy.

### *In Vitro* Cellular Internalization

Prior work has indicated that one of the advantages of a multifunctional multivalent approach is the ability to present an optimal ratio of bioactive for TPD, divergent from traditional chimeric strategies with 1:1 stoichiometry.^15, 23, 38-40^ Leveraging the tunability of PA self-assembly, 1:1, 2:1, and 3:1 CMA PA:α-Syn PA co-assemblies were prepared and assessed for their bioactivity. Assemblies containing more than 50 mol% α-Syn PA were not utilized in biological studies due to their inability to self-assemble into elongated nanofibers (Figure S4).

*In vitro* studies utilizing a human embryonic kidney cell line (HEK 293) were conducted to assess the cytotoxicity and cellular internalization capability of the control PA, the chimeric peptide, and bioactive PA co-assemblies (Figure 2). To assess cytotoxicity, a lactate dehydrogenase assay was performed on cells treated with up to 1 mM concentrations of the respective formulations for 48 hours. No significant cytotoxicity for all treatment concentrations and groups was observed compared to the no treatment controls up to 500 μM (Figure 2B).

**Figure 2.**
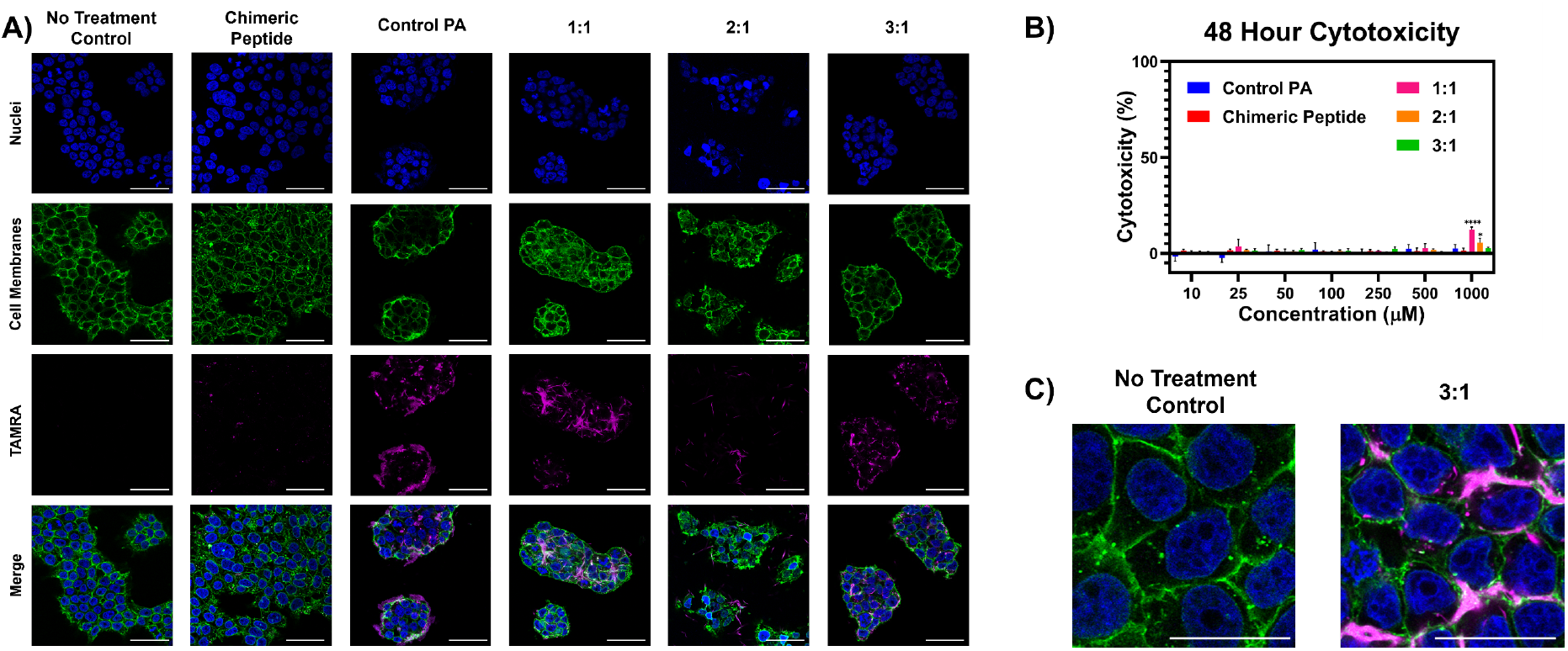
Biocompatibility and Cellular Internalization of Control and Bioactive PAs. A) Confocal microscopy images of HEK 293 cells fixed following 48-hour 100 µM treatments containing 5 mol% TAMRA-tagged compounds (magenta). Cell nuclei were stained with DAPI (blue), and cell membranes were stained with WGA 488 (green). Scale bars 50 µm. B) LDH cytotoxicity assay comparing the control PA, the chimeric peptide, and the TPD bioactive nanofibers 48 hours following incubations with HEK 293 cells. Data displayed as mean ± standard deviation, and a one-way analysis of variance (ANOVA) with a Tukey post hoc test was used to assess significance compared to a no treatment control (*p < 0.05; ****p<0.0001). C) Higher magnification merged images of no treatment control and 3:1. Scale bars are 20 µm.

Effective TPD is dependent on cellular internalization of the bioactive nanofibers to ultimately bind their intended intracellular protein targets. While many traditional therapeutics are either small enough to passively diffuse through the cell membrane or utilize an exogenous delivery vehicle, cellular internalization can be driven independently through several routes, one of which is the accumulation of cationic surface charge.^46, 47^ A common application of this mode is the incorporation of cell penetrating peptides with cationic sequences mimetic of the TAT sequence derived from HIV, with repeats of the amino acids arginine and lysine providing cationic charge.^12, 18, 28^ Both the CMA PA and the α-Syn PA present bioactive epitopes on the surface of the nanofibers with a net positive charge. While the net charges of the epitopes alone are unlikely as monomers to induce internalization, supramolecular co-assembly of these PAs produces nanofibers with a multivalent presentation of these sequences, creating a nanostructure with overall surface cationicity hypothesized to be suitable to facilitate cellular internalization.^47,48^ Cellular internalization was assessed qualitatively using confocal fluorescence microscopy of following 48-hour incubations of HEK 293 cells with 100 μM treatments of PAs co-assembled with 5 mol% of a control PA functionalized with 6-carboxytetramethylrhodamine (TAMRA, see supporting information for PA monomer structure and synthetic protocol) and of the chimeric peptide with a TAMRA-coupled N-terminus. Cells were fixed onto glass coverslips and stained for nuclei and cell membranes prior to imaging. Confocal microscopy indicated a high level of cellular internalization achieved by the bioactive TPD PA nanofibers at all co-assembly ratios compared to no treatment controls (Figure 2A and 2C). This was further supported by 3-dimensional images generated by confocal microscopy Z-stack reconstructions (Figure S5). Interestingly, the control PA also achieved comparable cellular internalization despite a largely anionic surface charge. We hypothesize this could be a contribution of the added cationicity from the incorporation of the TAMRA fluorescent label incorporated and the large surface area interaction with the cell membrane induced by the PA nanofiber architecture. Encouraged by high levels of cellular internalization of the bioactive TPD PA nanofibers, we next advanced to assessing α-Syn degradation in a model *in vitro* system.

### *In Vitro* Targeted Protein Degradation

To assess the ability of the bioactive PA nanofibers to induce the degradation of α-Syn, *in vitro* studies with a Förster resonance energy transfer (FRET) biosensor cell line in which HEK cells genetically modified to stably express fluorescently labeled α-Syn was used. The α-Syn in this biosensor cell line is present in two equal populations: one conjugated to a cyan fluorescent protein (CFP) and the other conjugated to yellow fluorescent protein (YFP).^49-51^ CFP and YFP are a FRET pair, allowing for qualitative observation of both reductions in α-Syn levels by monitoring overall fluorescent emission and aggregation by monitoring reductions in the YFP FRET emission.^52^ Biosensor HEK cells were grown to confluency and treated for 48 hours with either control PA nanofibers or one of our three co-assembled bioactive PA nanofibers. Cells were treated with 400 μM treatments, a concentration chosen to ideally induce an observable response while maintaining cell viability based on cytotoxicity results. Initially, to quantitatively assess α-Syn protein degradation, western blot experiments and densitometric analysis of protein bands were conducted (Figure 3). While all bioactive PA copolymers induced an observable average reduction in relative α-Syn levels, the 3:1 PA copolymer was the only formulation that exhibited statistically significant protein reduction, illustrating the importance of ligand stoichiometry in our multivalent nanofibers (see Figure S6 for additional western blots represented in Figure 3B). The 3:1 PA copolymer was down selected as our optimal formulation and utilized in further experiments.

**Figure 3.**
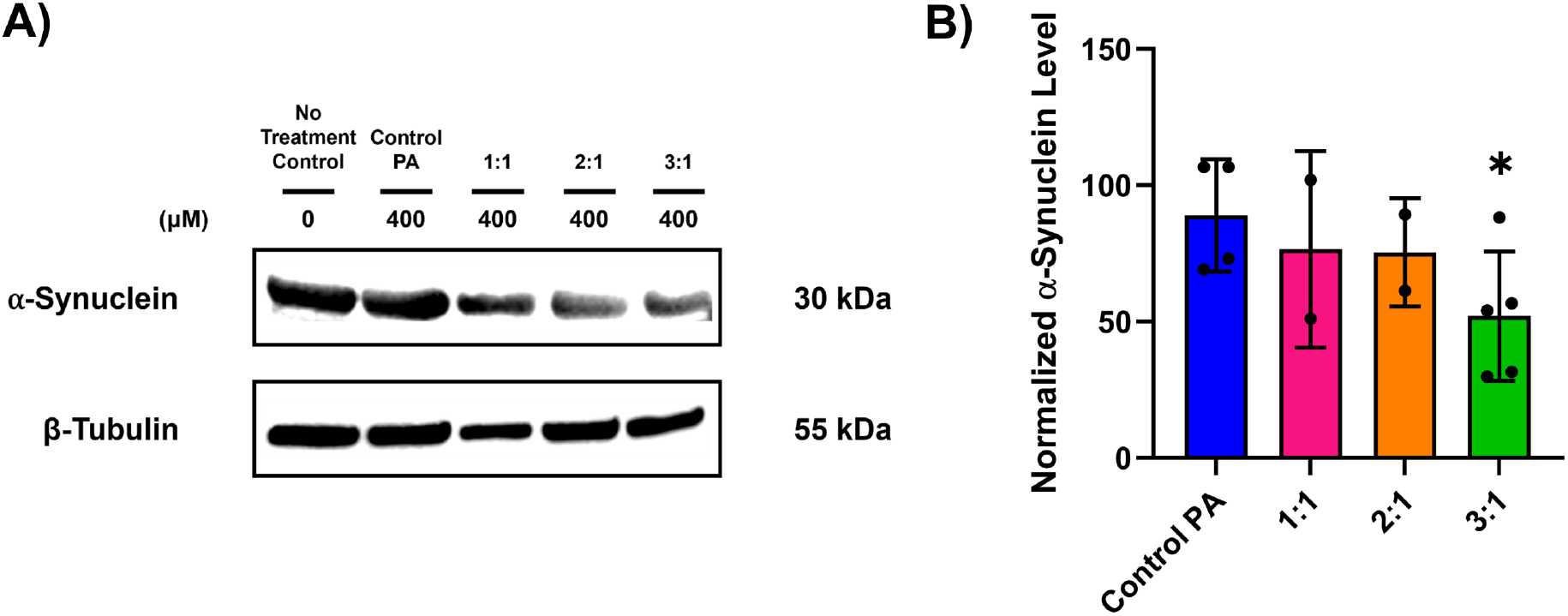
TPD via Application of PA supramolecular polymers. A) Representative protein bands observed via western blot analysis of cells incubated with 400 µM treatments with control PA and three different bioactive PA copolymers: 1:1, 2:1, and 3:1 (CMA PA: α-Syn PA, mol%) for 48 hours. B) Densitometric analysis of protein bands was conducted using ImageJ software. B-tubulin was used as a housekeeping control protein, and α-Syn levels were normalized to a no treatment control levels equaling 100% for each independent blot (n ≥ 2 for each treatment). Data displayed as mean ± standard deviation. Significance was assessed using a two-sided t-test with a hypothetical value of 100% (*p < 0.05).

To probe concentration dependence, TPD experiments were repeated with the 3:1 bioactive PA nanofibers, the control PA, and the chimeric peptide, comparing the efficacy of a macromolecular construct to that of a comparable linear peptide. Biosensor HEK cells were incubated with 100 and 400 μM concentrations of each treatment, respectively, followed by fixation and confocal microscopy (Figure 4A). Utilizing selective excitation of the CFP FRET donor and observance solely of the YFP FRET acceptor emission, representative images qualitatively indicated intracellular α-Syn reductions following only bioactive TPD PA nanofiber treatment, with an observable reduction in the intense puncta indicative of α-Syn intracellular aggregates (Figure 4A and 4B).^49, 50^ Quantitatively, western blot analysis demonstrated a dose-dependent reduction in α-Syn levels for cells treated with the bioactive TPD PA nanofibers, reducing intracellular α-Syn by on average 52% and up to approximately 75% compared to no treatment controls (Figure 4C and 4D) (see Figure S6 for additional western blots represented in Figure 4D). Further, confocal microscopy and western blot experiments indicate that the chimeric peptide and the control PA nanofibers do not effectively induce TPD, supporting the significance of the combined impact of the multivalent presentation nanofiber surface presentation being integral to enhancing TPD.

**Figure 4.**
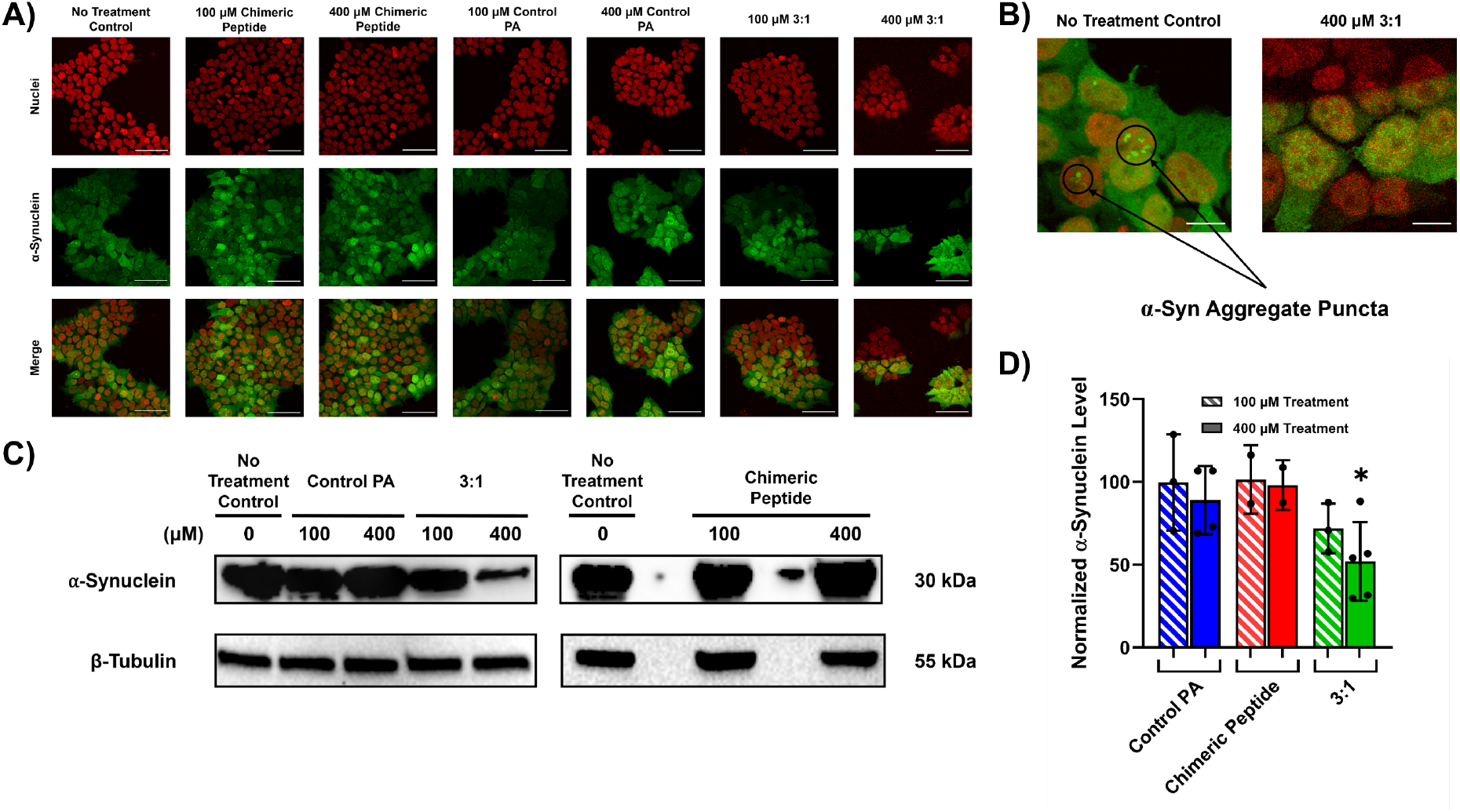
Dose-Dependent TPD of α-Syn. A) Confocal microscopy imaging of α-Syn HEK biosensor cells following 48-hour incubation with treatments. α-Syn indicated by YFP FRET emission (green), nuclei stained using NucSpot 650 (red). Scale bars are 50 µm. B) Higher magnification merged images of no treatment control and 400 µM 3:1. Scale bars are 10 µm. C) Representative protein bands observed via western blot analysis of treated cells. D) Densitometric analysis of protein bands observed in western blot was conducted using ImageJ software. B-tubulin was used as a housekeeping control protein, and α-Syn levels were normalized to a no treatment control levels equaling 100% for each independent blot (n ≥ 2 for each treatment). Data displayed as mean ± standard deviation. Significance was assessed using a two-sided t-test with a hypothetical value of 100% (*p < 0.05).

Following validation of the ability of our materials to induce the TPD of α-Syn, we moved to assess if protein degradation is occurring by the mechanism in which we hypothesized. CMA is a lysosome-dependent degradation pathway and should only occur in cells with functional lysosomes. To assess this lysosome-dependent degradation, 400 μM 3:1 bioactive PA nanofiber treatments were repeated with biosensor cells simultaneously treated with either the proteosome inhibitor N-benzyloxycarbonyl-L-leucyl-L-leucyl-L-leucinal (MG-132) or the lysosome inhibitor hydroxycholorquine (HCQ) (Figure 5).^11, 15, 23, 38^ We hypothesized that when co-treated with MG-132, α-Syn levels should still decrease upon treatment relative to control cells, but co-treatment with HCQ should render α-Syn levels largely unchanged relative to control cells. Following all treatments, relative α-Syn levels were increased compared to no treatment controls, which we attribute to impairment of basal intracellular α-Syn turnover caused by either inhibitor. Considering the large increase in α-Syn levels incubated with MG-132 compared to those incubated with HCQ, we hypothesize that most of the natural α-Syn turnover in these biosensor cells is proteosome-dependent. We believe that this makes it challenging to overcome proteosome inhibition and decrease α-Syn levels relative to no treatment controls in this aggressive, α-Syn producing model. However, we do observe a relative decrease in α-Syn levels in cells co-treated with 400 μM 3:1 and MG-132 compared to cells treated with MG-132 alone. On the contrary, α-Syn levels in cells co-treated with 400 μM 3:1 and HCQ were nearly identical to those treated with HCQ alone. Taken together, this supports our hypothesized lysosome-dependent CMA mechanism of degradation, demonstrating the capability of these multivalent nanofibers to provide a selective and predictable route to target POI elimination.

**Figure 5.**
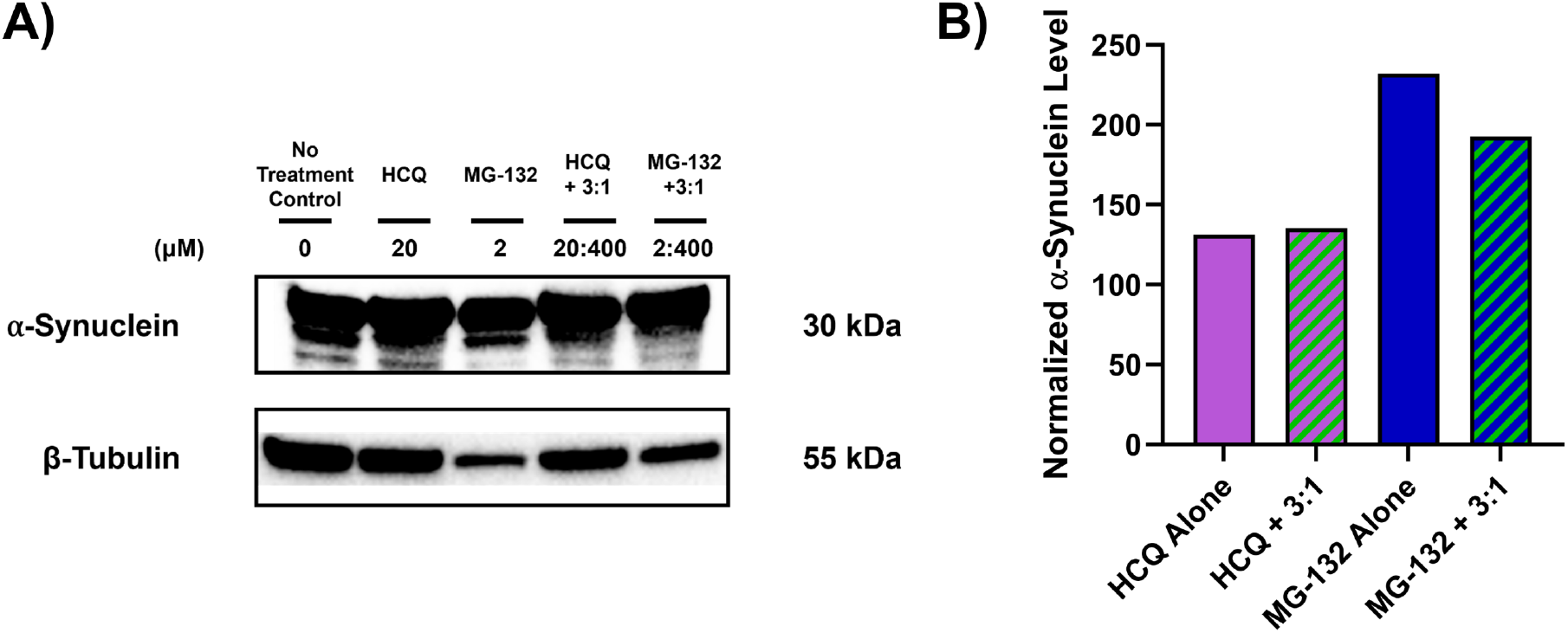
Mechanism of TPD. A) Representative protein bands observed via western blot analysis of cells incubated with 400 µM treatments with of the 3:1 bioactive PA copolymer co-incubated with MG-132 or HCQ. B) Densitometric analysis of protein bands was conducted using ImageJ software. B-tubulin was used as a housekeeping control protein, and α-Syn levels were normalized to a no treatment control levels equaling 100%.

## Conclusions

While TPD has been explored in a wide variety of small molecule and chimeric systems, macromolecular TPD approaches are only recently garnering attention, and unique targets like α-Syn are scarcely investigated. In this proof of principle study targeting α-Syn, inducing complete protein clearance was impeded by the competition between HEK cell doubling time and the time required for the treatments to achieve TPD, as well as the continuous overexpression of α-Syn. Despite the aggressiveness of this model, dose-dependent degradation of α-Syn was exhibited via application of multivalent nanofibers whose performance outcompeted that of a traditional, chimeric peptide exhibiting the same targeting epitopes. This underscores the potential for these PA nanofibers to further excel in primarily post-mitotic neuronal cells with relatively stable baseline protein concentrations, which is a focus of future studies.^11, 53^ Herein, we have demonstrated the promise of bioactive PA nanofibers as a dynamic and adaptable material platform poised to accelerate the design of next generation TPD therapeutics. The modular design of PAs enables precise targeting of a range of POIs, and the ease of modifying multivalent ligand display allows for POI-specific optimization, positioning PA nanofibers to serve as a universal platform through which efficient TPD can be achieved.

## Acknowledgements

Peptide synthesis and characterization in this work was supported by the BioPACIFIC Materials Innovation Platform of the National Science Foundation (NSF) under Award number DMR-1933487. Additional peptide synthesis in this study was also performed at the Peptide Synthesis Core Facility of the Simpson Querrey Institute for BioNanotechnology at Northwestern University. This facility has current support from the Soft and Hybrid Nanotechnology Experimental (SHyNE) Resource (NSF ECCS-2025633). Cell culture work included in this study was supported by the Mississippi INBRE, funded by an Institutional Development Award (IDeA) from the National Institute of General Medical Sciences of the NIH under award number P20GM103476. Additionally, the mass spectrometry facilities, and the transmission electron microscopy facilities utilized in this study were acquired through funding support from the National Science Foundation (NSF) Major Research Instrumentation (MRI) program under award number 2319932 and award number 2320081, respectively. The authors thank Mark Seniw for his design of Cinema 4D schematic drawings in the manuscript. Finally, the authors would like to acknowledge Dr. Penelope Jankoski for her support of this work and helpful guidance.

## FUNDING SOURCES

This work was primarily funded by the National Institutes of Health (NIH) National Institute of Biomedical Imaging and Bioengineering through award number R21EB033533 and R03EB033704. WSS gratefully acknowledges fellowship support from the Mississippi Space Grant Consortium (MSSGC) funded by the National Aeronautics and Space Administration (NASA) office of STEM engagement. TDC and AMD gratefully acknowledge support from the Arnold and Mabel Beckman Foundation.

### Notes

The authors declare the following competing financial interest(s): The authors WSS and TDC have filed a US provisional patent on aspects of the work described in this study.

## References

(1) Tang, H.; Andrikopoulos, N.; Li, Y.; Ke, S.; Sun, Y.; Ding, F.; Ke, P. C. Emerging biophysical origins and pathogenic implications of amyloid oligomers. Nature Communications 2025, 16 (1), 2937. DOI: 10.1038/s41467-025-58335-y.

(2) Tomoshige, S.; Ishikawa, M. PROTACs and Other Chemical Protein Degradation Technologies for the Treatment of Neurodegenerative Disorders. Angew Chem Int Ed Engl 2021, 60 (7), 3346–3354. DOI: 10.1002/anie.202004746From NLM.

(3) Jagaran, K.; Singh, M. Nanomedicine for Neurodegenerative Disorders: Focus on Alzheimer’s and Parkinson’s Diseases. International Journal of Molecular Sciences 2021, 22 (16), 9082.

(4) Poovaiah, N.; Davoudi, Z.; Peng, H.; Schlichtmann, B.; Mallapragada, S.; Narasimhan, B.; Wang, Q. Treatment of neurodegenerative disorders through the blood–brain barrier using nanocarriers. Nanoscale 2018, 10 (36), 16962–16983, 10.1039/C8NR04073G. DOI: 10.1039/C8NR04073G.

(5) Fang, Y.; Wang, J.; Zhao, M.; Zheng, Q.; Ren, C.; Wang, Y.; Zhang, J. Progress and Challenges in Targeted Protein Degradation for Neurodegenerative Disease Therapy. Journal of Medicinal Chemistry 2022, 65 (17), 11454–11477. DOI: 10.1021/acs.jmedchem.2c00844.

(6) Xie, X.; Yu, T.; Li, X.; Zhang, N.; Foster, L. J.; Peng, C.; Huang, W.; He, G. Recent advances in targeting the “undruggable” proteins: from drug discovery to clinical trials. Signal Transduction and Targeted Therapy 2023, 8 (1), 335. DOI: 10.1038/s41392-023-01589-z.

(7) Coleman, N.; Rodon, J. Taking Aim at the Undruggable. American Society of Clinical Oncology Educational Book 2021, (41), e145–e152. DOI: 10.1200/EDBK_325885 (accessed 2024/02/22).

(8) Wen, T.; Chen, J.; Zhang, W.; Pang, J. Design, Synthesis and Biological Evaluation of α-Synuclein Proteolysis-Targeting Chimeras. Molecules 2023, 28 (11), 4458.

(9) Fields, C. R.; Bengoa-Vergniory, N.; Wade-Martins, R. Targeting Alpha-Synuclein as a Therapy for Parkinson’s Disease. Frontiers in Molecular Neuroscience 2019, 12, Review. DOI: 10.3389/fnmol.2019.00299.

(10) Shaltiel-Karyo, R.; Frenkel-Pinter, M.; Egoz-Matia, N.; Frydman-Marom, A.; Shalev, D. E.; Segal, D.; Gazit, E. Inhibiting α-synuclein oligomerization by stable cell-penetrating β-synuclein fragments recovers phenotype of Parkinson’s disease model flies. PLoS One 2010, 5 (11), e13863. From NLM.

(11) Lee, J.; Sung, K. W.; Bae, E.-J.; Yoon, D.; Kim, D.; Lee, J. S.; Park, D.-h.; Park, D. Y.; Mun, S. R.; Kwon, S. C.; et al. Targeted degradation of α-synuclein aggregates in Parkinson’s disease using the AUTOTAC technology. Molecular Neurodegeneration 2023, 18 (1), 41. DOI: 10.1186/s13024-023-00630-7.

(12) Jin, J. W.; Fan, X.; Del Cid-Pellitero, E.; Liu, X. X.; Zhou, L.; Dai, C.; Gibbs, E.; He, W.; Li, H.; Wu, X.; et al. Development of an α-synuclein knockdown peptide and evaluation of its efficacy in Parkinson’s disease models. Commun Biol 2021, 4 (1), 232. DOI: 10.1038/s42003-021-01746-6From NLM.

(13) Amirian, R.; Badrbani, M. A.; Derakhshankhah, H.; Izadi, Z.; Shahbazi, M.-A. Targeted protein degradation for the treatment of Parkinson’s disease: Advances and future perspective. Biomedicine & Pharmacotherapy 2023, 166, 115408. DOI:10.1016/j.biopha.2023.115408.

(14) Burslem, G. M.; Crews, C. M. Proteolysis-Targeting Chimeras as Therapeutics and Tools for Biological Discovery. Cell 2020, 181 (1), 102–114. From NLM.

(15) Wang, J.; Wang, Y.; Yang, F.; Luo, Q.; Hou, Z.; Xing, Y.; Lu, F.; Li, Z.; Yin, F. A Novel Lysosome Targeting Chimera for Targeted Protein Degradation via Split-and-Mix Strategy. Acs Chem Biol 2024, 19 (5), 1161–1168. DOI: 10.1021/acschembio.4c00092.

(16) Wang, H.; Zhou, R.; Xu, F.; Yang, K.; Zheng, L.; Zhao, P.; Shi, G.; Dai, L.; Xu, C.; Yu, L.; et al. Beyond canonical PROTAC: biological targeted protein degradation (bioTPD). Biomater Res 2023, 27 (1), 72. DOI: 10.1186/s40824-023-00385-8From NLM.

(17) Bondeson, D. P.; Mares, A.; Smith, I. E. D.; Ko, E.; Campos, S.; Miah, A. H.; Mulholland, K. E.; Routly, N.; Buckley, D. L.; Gustafson, J. L.; et al. Catalytic in vivo protein knockdown by small-molecule PROTACs. Nature Chemical Biology 2015, 11 (8), 611–617. DOI: 10.1038/nchembio.1858.

(18) Fan, X.; Jin, W. Y.; Lu, J.; Wang, J.; Wang, Y. T. Rapid and reversible knockdown of endogenous proteins by peptide-directed lysosomal degradation. Nat Neurosci 2014, 17 (3), 471–480. From NLM.

(19) Kaushik, S.; Cuervo, A. M. The coming of age of chaperone-mediated autophagy. Nat Rev Mol Cell Biol 2018, 19 (6), 365–381. From NLM.

(20) Valdor, R.; Martinez-Vicente, M. The Role of Chaperone-Mediated Autophagy in Tissue Homeostasis and Disease Pathogenesis. Biomedicines 2024, 12 (2). From NLM.

(21) Shao, J.; Lin, X.; Wang, H.; Zhao, C.; Yao, S. Q.; Ge, J.; Zeng, S.; Qian, L. Targeted Degradation of Cell-Surface Proteins via Chaperone-Mediated Autophagy by Using Peptide-Conjugated Antibodies. Angew Chem Int Ed Engl 2024, 63 (18), e202319232. DOI: 10.1002/anie.202319232From NLM.

(22) Bauer, P. O.; Goswami, A.; Wong, H. K.; Okuno, M.; Kurosawa, M.; Yamada, M.; Miyazaki, H.; Matsumoto, G.; Kino, Y.; Nagai, Y.; et al. Harnessing chaperone-mediated autophagy for the selective degradation of mutant huntingtin protein. Nat Biotechnol 2010, 28 (3), 256–263. DOI: 10.1038/nbt.1608From NLM.

(23) Song, C.; Jiao, Z.; Hou, Z.; Wang, R.; Lian, C.; Xing, Y.; Luo, Q.; An, Y.; Yang, F.; Wang, Y.; et al. Selective Protein of Interest Degradation through the Split-and-Mix Liposome Proteolysis Targeting Chimera Approach. Journal of the American Chemical Society 2023, 145 (40), 21860–21870. DOI: 10.1021/jacs.3c05948.

(24) Wang, Y.; Zhao, J.; Yuan, H.; Li, S. Advances in Peptide-Based Chimeric Strategies for Targeted Protein Degradation. Journal of Medicinal Chemistry 2025, 68 (24), 25708–25728. DOI: 10.1021/acs.jmedchem.5c02379.

(25) Borsari, C.; Trader, D. J.; Tait, A.; Costi, M. P. Designing Chimeric Molecules for Drug Discovery by Leveraging Chemical Biology. J Med Chem 2020, 63 (5), 1908–1928. DOI: 10.1021/acs.jmedchem.9b01456 From NLM.

(26) Sun, H.; Qiao, B.; Choi, W.; Hampu, N.; McCallum, N. C.; Thompson, M. P.; Oktawiec, J.; Weigand, S.; Ebrahim, O. M.; de la Cruz, M. O.; et al. Origin of Proteolytic Stability of Peptide-Brush Polymers as Globular Proteomimetics. ACS Central Science 2021, 7 (12), 2063–2072. DOI: 10.1021/acscentsci.1c01149.

(27) Wang, L.; Wang, N.; Zhang, W.; Cheng, X.; Yan, Z.; Shao, G.; Wang, X.; Wang, R.; Fu, C. Therapeutic peptides: current applications and future directions. Signal Transduction and Targeted Therapy 2022, 7 (1), 48. DOI: 10.1038/s41392-022-00904-4.

(28) Tong, Y.; Zhu, W.; Chen, J.; Zhang, W.; Xu, F.; Pang, J. Targeted Degradation of Alpha-Synuclein by Autophagosome-Anchoring Chimera Peptides. Journal of Medicinal Chemistry 2023, 66 (17), 12614–12628. DOI: 10.1021/acs.jmedchem.3c01303.

(29) da Silva, R. M. P.; van der Zwaag, D.; Albertazzi, L.; Lee, S. S.; Meijer, E. W.; Stupp, S. I. Super-resolution microscopy reveals structural diversity in molecular exchange among peptide amphiphile nanofibres. Nature Communications 2016, 7 (1), 11561. DOI: 10.1038/ncomms11561.

(30) Cui, H.; Webber, M. J.; Stupp, S. I. Self-assembly of peptide amphiphiles: from molecules to nanostructures to biomaterials. Biopolymers 2010, 94 (1), 1–18. DOI: 10.1002/bip.21328 From NLM.

(31) Yuan, S. C.; Lewis, J. A.; Sai, H.; Weigand, S. J.; Palmer, L. C.; Stupp, S. I. Peptide Sequence Determines Structural Sensitivity to Supramolecular Polymerization Pathways and Bioactivity. J Am Chem Soc 2022, 144 (36), 16512–16523. DOI: 10.1021/jacs.2c05759From NLM.

(32) Sangji, M. H.; Sai, H.; Chin, S. M.; Lee, S. R.; I, R. S.; Palmer, L. C.; Stupp, S. I. Supramolecular Interactions and Morphology of Self-Assembling Peptide Amphiphile Nanostructures. Nano Lett 2021, 21 (14), 6146–6155. DOI: 10.1021/acs.nanolett.1c01737 From NLM.

(33) Edelbrock, A. N.; Clemons, T. D.; Chin, S. M.; Roan, J. J. W.; Bruckner, E. P.; Álvarez, Z.; Edelbrock, J. F.; Wek, K. S.; Stupp, S. I. Superstructured Biomaterials Formed by Exchange Dynamics and Host–Guest Interactions in Supramolecular Polymers. Advanced Science 2021, 8 (8), 2004042. DOI:10.1002/advs.202004042.

(34) Jankoski, P. E.; Wallace, Z. M.; DiMartino, L. R.; Shrestha, J.; Davis, A. M.; Owolabi, I.; Flynt, A. S.; Clemons, T. D. Combating Reactive Oxygen Species (ROS) with Antioxidant Supramolecular Polymers. ACS Applied Materials & Interfaces 2025, 17 (24), 35275–35287. DOI: 10.1021/acsami.5c06967.

(35) Lee, S. S.; Fyrner, T.; Chen, F.; Álvarez, Z.; Sleep, E.; Chun, D. S.; Weiner, J. A.; Cook, R. W.; Freshman, R. D.; Schallmo, M. S.; et al. Sulfated glycopeptide nanostructures for multipotent protein activation. Nature Nanotechnology 2017, 12 (8), 821–829. DOI: 10.1038/nnano.2017.109.

(36) Jankoski, P. E.; Masoud, A.-R.; Dennis, J.; Trinh, S.; DiMartino, L. R.; Shrestha, J.; Marrero, L.; Hobden, J.; Carter, J.; Schoen, J.; et al. Bioactive Supramolecular Polymers for Skin Regeneration Following Burn Injury. Biomacromolecules 2025, 26 (8), 5471–5482. DOI: 10.1021/acs.biomac.5c01107.

(37) Ledford, B. T.; Akerman, A. W.; Sun, K.; Gillis, D. C.; Weiss, J. M.; Vang, J.; Willcox, S.; Clemons, T. D.; Sai, H.; Qiu, R.; et al. Peptide Amphiphile Supramolecular Nanofibers Designed to Target Abdominal Aortic Aneurysms. ACS Nano 2022, 16 (5), 7309–7322. DOI: 10.1021/acsnano.1c06258.

(38) Song, C.; Jiao, Z.; Hou, Z.; Xing, Y.; Sha, X.; Wang, Y.; Chen, J.; Liu, S.; Li, Z.; Yin, F. Versatile Split-and-Mix Liposome PROTAC Platform for Efficient Degradation of Target Protein In Vivo. JACS Au 2024, 4 (8), 2915–2924. DOI: 10.1021/jacsau.4c00278.

(39) Zhan, M.-m.; Chen, H.; He, M.; Song, C.; Jiao, Z.; Liu, N.; Liu, Z.; Hou, Z.; Chen, Y.; Song, Z.; et al. Highly efficient and long-acting split-and-mix proteolysis targeting chimera based on self-assembled polylactic acid. Nature Communications 2025, 16 (1), 10555. DOI: 10.1038/s41467-025-65590-6.

(40) Yang, F.; Luo, Q.; Wang, Y.; Liang, H.; Wang, Y.; Hou, Z.; Wan, C.; Wang, Y.; Liu, Z.; Ye, Y.; et al. Targeted Biomolecule Regulation Platform: A Split-and-Mix PROTAC Approach. Journal of the American Chemical Society 2023, 145 (14), 7879–7887. DOI: 10.1021/jacs.2c12824.

(41) Wang, M. M.; Truica, M. I.; Gattis, B. S.; Oktawiec, J.; Sagar, V.; Basu, A. A.; Bertin, P. A.; Zhang, X.; Abdulkadir, S. A.; Gianneschi, N. C. Heterobifunctional proteomimetic polymers for targeted degradation of MYC and KRAS. Nature Communications 2026, 17 (1), 1706. DOI: 10.1038/s41467-026-68913-3.

(42) Sangji, M. H.; Lee, S. R.; Sai, H.; Weigand, S.; Palmer, L. C.; Stupp, S. I. Self-Sorting vs Coassembly in Peptide Amphiphile Supramolecular Nanostructures. ACS Nano 2024, 18 (24), 15878–15887. DOI: 10.1021/acsnano.4c03083.

(43) Edelbrock, A. N.; Clemons, T. D.; Chin, S. M.; Roan, J. J. W.; Bruckner, E. P.; Alvarez, Z.; Edelbrock, J.; Wek, K. S.; Stupp, S. I. Superstructured Biomaterials Formed by Exchange Dynamics and Host-Guest Interactions in Supramolecular Polymers. Advanced Science 2021, 2004042.

(44) Klein, M. K.; Kassam, H. A.; Lee, R. H.; Bergmeier, W.; Peters, E. B.; Gillis, D. C.; Dandurand, B. R.; Rouan, J. R.; Karver, M. R.; Struble, M. D.; et al. Development of Optimized Tissue-Factor-Targeted Peptide Amphiphile Nanofibers to Slow Noncompressible Torso Hemorrhage. ACS Nano 2020, 14 (6), 6649–6662. DOI: 10.1021/acsnano.9b09243.

(45) Jankoski, P. E.; Shrestha, J.; Swetman, W. S.; Livingston, H.; Sorrell, J.; Clemons, T. D. Cell-Laden Supramolecular and Covalent Polymer Hydrogels for High-Shear Delivery: A Design of Experiments Approach. Chemistry of Materials 2026. DOI: 10.1021/acs.chemmater.5c03073.

(46) Blum, A. P.; Yin, J.; Lin, H. H.; Oliver, B. A.; Kammeyer, J. K.; Thompson, M. P.; Gilson, M. K.; Gianneschi, N. C. Stimuli Induced Uptake of Protein-Like Peptide Brush Polymers. nChemistry 2022, 28 (5), e202103438. From NLM.

(47) Sousa de Almeida, M.; Susnik, E.; Drasler, B.; Taladriz-Blanco, P.; Petri-Fink, A.; Rothen-Rutishauser, B. Understanding nanoparticle endocytosis to improve targeting strategies in nanomedicine. Chemical Society Reviews 2021, 50 (9), 5397–5434, 10.1039/D0CS01127D. DOI: 10.1039/D0CS01127D.

(48) Mondal, M.; Swetman, W. S.; Karim, S.-U.; Shrestha, S.; Davis, A. M.; Bai, F.; Huang, F.; Clemons, T. D.; Rangachari, V. Disulfide cross-linked redox-sensitive peptide condensates are efficient cell delivery vehicles of molecular cargo. Proceedings of the National Academy of Sciences 2025, 122 (41), e2515427122. DOI: 10.1073/pnas.2515427122 (accessed 2026/01/15).

(49) Holmes, B. B.; Furman, J. L.; Mahan, T. E.; Yamasaki, T. R.; Mirbaha, H.; Eades, W. C.; Belaygorod, L.; Cairns, N. J.; Holtzman, D. M.; Diamond, M. I. Proteopathic tau seeding predicts tauopathy in vivo. Proceedings of the National Academy of Sciences 2014, 111 (41), E4376–E4385. DOI: 10.1073/pnas.1411649111 (accessed 2026/02/17).

(50) Yamasaki, T. R.; Holmes, B. B.; Furman, J. L.; Dhavale, D. D.; Su, B. W.; Song, E.-S.; Cairns, N. J.; Kotzbauer, P. T.; Diamond, M. I. Parkinson’s disease and multiple system atrophy have distinct α-synuclein seed characteristics. Journal of Biological Chemistry 2019, 294 (3), 1045–1058. DOI: 10.1074/jbc.RA118.004471 (accessed 2026/02/17).

(51) Furman, J. L.; Holmes, B. B.; Diamond, M. I. Sensitive Detection of Proteopathic Seeding Activity with FRET Flow Cytometry; 1940–087X, 2015. DOI: doi:10.3791/53205.

(52) Maina, K. N.; Smet-Nocca, C.; Bitan, G. Using FRET-Based Biosensor Cells to Study the Seeding Activity of Tau and α-Synuclein. Methods Mol Biol 2023, 2551, 125–145. From NLM.

(53) Breunig, Joshua J.; Haydar Tarik F.; Rakic, P. Neural Stem Cells: Historical Perspective and Future Prospects. Neuron 2011, 70 (4), 614–625. DOI:10.1016/j.neuron.2011.05.005.

