## Supplementary Information for "Leveraging Supramolecular Polymers to Induce the Targeted Protein Degradation of α-Synuclein"

#### Supporting Information

**2.4.5 Western Blot Analysis** ... 6
**3. Supplemental Figures** ... 7
**Figure S1. Electrospray ionization mass spectrometry analysis of purified peptides.** ... 7
**Figure S2. Critical aggregate concentration determination by Nile red assay** ... 8
**Figure S3. Nanostructure Characterization of Chimeric Peptide.** ... 8
**Figure S4. TEM Imaging of 1:3 (CMA Pa, α-Syn PA, mol%) co-assembly.** ... 9
**Figure S5. Confocal microscopy of PA cellular internalization Z-stacks.** ... 9
**Figure S6. Additional western blot experiments performed followed TPD treatments.** ... 9

### 1. Materials and Methods

#### 1.1 Materials

Rink amide Polystyrene Resin, Fmoc protected amino acids, and ethyl cyanoglyoxlate-2oxime ( Oxyma) were purchased from CEM peptides. Diethyl ether, trifluoroacetic acid (TFA), N-dimethylformamide (DMF), acetonitrile, diisopropylcarbodiimide (DIC), triisopropylsilane (TIS), ethane-1,2-dithiol (EDT), Uranyl Acetate and all other solvents were purchased from ThermoFisher Scientific (USA) or Sigma Aldrich Corporation (USA) at the highest purity. Dulbecco's modified eagle medium (DMEM), heat inactivated fetal bovine serum (FBS), and penicillin-streptomycin were all purchased from ThermoFisher, sterile, and tissue treated 96- and 6-well plates were obtained from CellTreat (USA). The CyQUANT Lactate Dehydrogenase (LDH) Assay was purchased from ThermoFisher (USA). Hexafluoroisopropanol (HFIP) was obtained from AA blocks and Nile Red was purchased from ApexBio.

#### 1.2 Peptide Synthesis

Peptides were synthesized on a Liberty Blue 2.0 automated peptide synthesizer (CEM) through standard 9-fluorenyl methoxycarbonyl (Fmoc)- based solid phase peptide synthesis. Peptide synthesis was performed at 0.25 mmol scale using Rink Amide Polystyrene Resin (0.65 mmol/g loading, 100-200 mesh). Deprotection of Fmoc protecting groups was carried out using 20 v/v% piperidine in DMF. Each amino acid addition was carried out using Fmoc-protected amino acids (0.2 M), DIC (1M), and Oxyma (1 M) in DMF. After the final Fmoc deprotection, the resin beads were washed 3x using DMF. The peptide then underwent global deprotection and cleavage from the resin beads through gentle shaking in TFA/TIS/$H_2O$/EDT (92.5: 2.5:2.5:2.5) cleavage cocktail for 3 hours at room temperature. Peptides were then precipitated in cold diethyl ether and collected via centrifugation. The peptide pellet was then resuspended in diethyl ether and chilled for four hours. It was recentrifuged and the diethyl ether supernatant was decanted from the peptide pellet, which in turn was allowed to air dry. Crude peptides were purified on a Prodigy preparative reverse-phase HPLC (CEM) with a water/acetonitrile gradient (containing 0.1% $NH_4OH$ or 0.1% TFA). The mass and identity of the eluting fractions containing the desired peptides were confirmed using electrospray ionization (ESI)- mass spectrometry (MS) on a Thermo Scientific Orbitrap Exploris™ 240. Purity was confirmed with a demonstrated purity of greater than 95%.

1.2.1 Control PA Synthesis

The following peptide sequence $C_{16}V_2A_2E_2$ was synthesized on Rink amide MBHA resin making use of the CEM Liberty microwave-assisted peptide synthesizer and protocols described above in section 1.1.

1.2.2 CMA Synthesis

The following peptide sequence $C_{16}V_2A_2E_2$-$G_2$-KFERQ was synthesized on Rink amide MBHA resin making use of the CEM Liberty microwave-assisted peptide synthesizer and protocols described above in section 1.1.

1.2.2 α-Syn PA Synthesis

The following peptide sequence $C_{16}V_2A_2E_4$-GVLYVGSKTR was synthesized on Rink amide MBHA resin making use of the CEM Liberty microwave-assisted peptide synthesizer and protocols described above in section 1.1.

1.2.2 Chimeric Peptide Synthesis

The following peptide sequence GVLYVGSKTR-GSGS-KFERQ was synthesized on Rink amide MBHA resin making use of the CEM Liberty microwave-assisted peptide synthesizer and protocols described above in section 1.1.

1.2.3 TAMRA Control PA Synthesis

The following peptide sequence $C_{16}V_2A_2E_2$-K(Mtt) was synthesized on Rink amide MBHA resin making use of the CEM Liberty microwave-assisted peptide synthesizer and protocols described above in section 1.1. The terminal lysine $\varepsilon$-amine, protected with 4-methyltrityl (Mtt), was selectively deprotected on resin through the addition of a deprotection cocktail 3:5:92 TFA/TIPS/DCM, for multiple 5 min washes until yellow color was no longer seen in solution. Successful deprotection and subsequent coupling was verified through ninhydrin colorimetric assay (Kaiser test). Carboxytetramethyl rhodamine (TAMRA) (2 equiv) was coupled to the free $\varepsilon$-amine with 2 equiv TAMRA, 2 equiv PyBOP and 6 equiv DIEA in DMF for 16 h on a mechanical peptide shaker. Following successful coupling, the PA was cleaved, purified and stored as described in section 1.2.1.

1.2.3 TAMRA Chimeric Peptide Synthesis

Chimeric peptide was synthesized as described above in section 1.2.2. Prior to global deprotection and resin cleavage TAMRA (2 equiv) was coupled to the free amine on the N-terminus of the peptide with 2 equiv TAMRA, 2 equiv PyBOP and 6 equiv DIEA in DMF for 16 h on a mechanical peptide shaker. Following successful coupling, the PA was cleaved, purified and stored as described in section 1.2.1.

##### 1.3.1 Peptide Amphiphile Nanofiber Preparation

Lyophilized pure peptides yielded from synthesis were dissolved in 100 mM HCl and lyophilized again to neutralize any lingering TFA. The PAs were then weighed out into Eppendorf tubes to be at 10 mM concentration with a working volume of 1 mL. The PAs were then dissolved in milli-Q water and slowly pH adjusted to a pH between 7 and 8 using 1 M NaOH being careful not to overshoot. The PAs were then lyophilized and resuspended in working volume to obtain 10 mM stock solutions. To prepare the PA coassemblies, appropriate volumes of 10 mM CMA PA and α-Syn PA were mixed. For instance, 750 µL of CMA PA and 250 µL of α-Syn PA were mixed for the 3:1 blend. The solutions were then lyophilized. The dry powders were dissolved in HFIP and left to evaporate overnight and then redissolved in 1 mL of DI water and lyophilized. PAs and the chimeric peptide were stored at -20 °C until use. Following resuspension in milli-Q water to obtain 10 mM working stock solutions, PAs were thermally annealed at 80 °C for 30 minutes and slowly cooled at 1 °C per minute back to room temperature.

#### 2. Material Characterization

##### 2.1 Nile Red Assay

Stock solutions of Nile Red were prepared at 10 mM in DMSO and subsequently diluted with deionized (DI) water to achieve a final working concentration of 100 µM. PA samples were diluted in DI water to produce a concentration range of 0 to 500 µM. In a 96-well plate, 90 µL of PA solution and 10 µL of Nile Red solution were added to each well, mixed thoroughly, and incubated at room temperature for 3 hours, with intermittent tapping to promote incorporation. Each condition was performed in triplicate. After incubation, samples were analyzed using a Biotek Synergy H1 Microplate Reader (Agilent), with excitation set to 550 nm and emission measured in 2 nm increments from 580 to 720 nm. The mean maximum relative fluorescence units (RFU) were plotted against the logarithm of the concentration, and the critical aggregation concentration (CAC) was determined as the intersection point of the curves corresponding to the absence and presence of fluorescence.

##### 2.2 Circular Dichroism (CD)

PAs were prepared as described above and diluted to 100 – 500 µM in milli-q water. CD spectra was recorded in a 1 mm pathlength cuvette on a J-815 (Jasco, Easton, MD) spectropolarimeter. Continuous scanning mode was used with a scanning speed of 100 nm per minute over a measurement range of 190 – 300 nm. The high-tension voltage (HT) was also monitored to ensure that the measurement was not saturated. Three measurements were obtained, and the buffer sample was run as a background that was subtracted.

##### 2.3 Transmission Electron Microscopy (TEM)

200-mesh copper grids (Ted Pella) were used as purchased. PA solution was diluted to 0.5 mM and 5 µL was dropped on the grid and left to sit for 5 minutes. Excess solution was wicked away and a drop of uranyl acetate was added as a stain to sit for 2 minutes. Excess solution was wicked away and 5 µL of deionized water was added and left to sit for 5 minutes. The solution was wicked away and grids were left to air dry prior to visualization using a JEOL JEM120i instrument at 120 kV.

#### 2.4 Cell Work

##### 2.4.1 Cell Culture Maintenance

Human embryonic kidney cells (HEK 293) and α-Syn Biosensor HEK cells were maintained in Dulbecco's modified eagle medium (DMEM), supplemented with 10% fetal bovine serum (FBS) and 1% penicillin-streptomycin. Cells were cultured at 37° C, 5% $CO_2$ in tissue treated flasks and used at confluence of ~ 80%.

##### 2.4.2 Cytotoxicity

HEK 293 cells were seeded in a 96 well plate (5 x $10^4$ cells/mL, 100 µL volume per well). Seeded cells were incubated at 37 °C and 5% $CO_2$ overnight to allow cells to adhere. Following adherence, PA solution was added to the media to incubate, and each concentration was performed in triplicate. Nuclease free water was used as a spontaneous control, and Triton X-100 was used as a positive control for 100 % cytotoxicity (i.e. complete LDH release). Plates were incubated for 48 hours at 37 °C and 5% $CO_2$, before collecting media to assess LDH release with the CyQUANT LDH assay following the manufacturers protocols. A microplate reader was used to assess the absorbance at 490 nm with a reference wavelength of 690 nm. % Cytotoxicity was calculated using equations 1 and 2 below.

$$(1)\ LDH\ Activity = A_{sample,490} - A_{sample,680}$$

$$(2)\ \%\ Cytotoxicity\ \left(\frac{LDH\ Activity_{sample} - LDH\ Activity_{spontaneous}}{LDH\ Activity_{Max\ Lysis} - LDH\ Activity_{spontaneous}}\right) * 100$$

##### 2.4.3 Treatment Incubations for Confocal Imaging and Western Blot Analysis

Cells were seeded in 6 well plates following the protocol described below in section 2.4.4. To prepare treatment media, 10 mM PA and chimeric peptide stock solutions were stepwise diluted using unsupplemented DMEM to final desired treatment concentrations. To treat cells, maintenance cell culture media was removed from a given well, cells were washed three times with PBS, and, wells were refilled with treatment media to the normal maintenance volume of 2.0 mL per well. After four hours, 2.0 mL of DMEM supplemented with 2x concentrations of normal maintenance cell culture media (20% FBS and 2% penicillin-streptomycin) was added to each well, bringing each well's total volume to 4.0 mL and supplement concentrations to normal maintenance levels. 48 hours after initial treatment incubation, cells were fixed for confocal imaging or harvested for western blot analysis following the protocols described below in sections 2.4.4 and 2.4.5. For western blot studies in which cells were treated with the inhibitors HCQ or MG-132, cells were pre-treated with the inhibitors (final concentration 20 µM HCQ and 2 µM MG-132) two hours prior to TPD experiment start in normal maintenance cell culture media and further incubated with cells for the entire duration of a TPD experiment following the same stepwise dilution protocol outlined above.

##### 2.4.4 Fixed Cell Confocal Imaging

Glass cover slips (18 x 18 mm, No. 1.5 glass) were washed (60% MeOH, 30% milli-q water, 10% NaOH) three times, rinsed three times with milli-q water, and sterilized with UV light. Following this preparation, cover slips were set in a 6 well plate and coated with poly-L-lysine (2 hours at 37 °C). Cover slips were rinsed three times with PBS prior to seeding cells. Using HEK 293 cells for cellular internalization and α-Syn Biosensor HEK cells for TPD studies, cells were seeded on the prepared glass cover slips in the 6 well plate (1 x $10^5$ cells/mL, 2.0 mL volume per well). Seeded cells were incubated at 37 °C and 5% $CO_2$ overnight to allow cells to adhere. Following completion of an experiment, cells were rinsed three times with PBS fixed via incubation in 4 w/v% paraformaldehyde (PFA) for 15 minutes at room temperature. Cells were rinsed three times with PBS to remove excess PFA. For cellular internalization studies, cell membranes were stained with WGA 488 (Biotium) according to the manufacturer's instructions, and cell nuclei were stained with DAPI Fluoromount-G Mounting Medium (Southern Biotech) according to the manufacturer's instructions. Z-stack reconstructions were taken to 3D extra- and intracellular localization of PAs incubated with cells. For TPD studies, cell nuclei were stained with NucSpot 650 (Biotium) according to the manufacturer's instructions. Glass coverslips were mounted on glass slides and sealed prior to imaging. Fixed cells were imaged on a Leica STELLARIS STED Super-Resolution Confocal Microscope.

###### 2.4.5 Western Blot Analysis

Treated cells were lysed, and proteins were extracted using the Thermo Scientific NE-PER Nuclear and Cytoplasmic Extraction Kit according to the manufacturer's instructions. Aliquots of the protein samples were mixed with 4× Laemmli sample buffer, heated at 100°C for 5 minutes, and loaded onto the SDS-PAGE setup. Equal amounts of protein (roughly calculated based on transfection efficiency) were loaded onto Mini-PROTEAN® TGX 4–20% precast SDS-PAGE gels (BioRad) and were electrophoresed. The gels were then transferred onto a 0.45 µm nitrocellulose membrane (Amersham Protran Premium, GE Life Sciences). Following transfer, membranes were blocked for 1 h at room temperature in blocking buffer containing 5% non-fat dry milk and 0.1% Tween-20 in Tris-buffered saline (TBS). Membranes were incubated overnight at 4 °C with primary antibodies against α-synuclein (Millipore) and β-tubulin (Cell Signaling Technology), diluted in blocking buffer. Membranes were then washed three times (5 minutes each) with Tris-buffered saline with Tween 20 (TBST) buffer and incubated for an additional 1 hour at room temperature with the appropriate horseradish peroxidase (HRP)-conjugated secondary antibodies (while HRP-conjugated anti-rabbit IgG was used for β-tubulin, HRP-conjugated anti-mouse IgG was used for α-synuclein). Membranes were then washed three additional times with TBST for 5 minutes each. Protein bands were then visualized using SuperSignal™ West Pico PLUS Chemiluminescent Substrate (Thermo Fisher Scientific) and imaged using a GelDoc Molecular Imager (Bio-Rad).

#### 3. Supplemental Figures

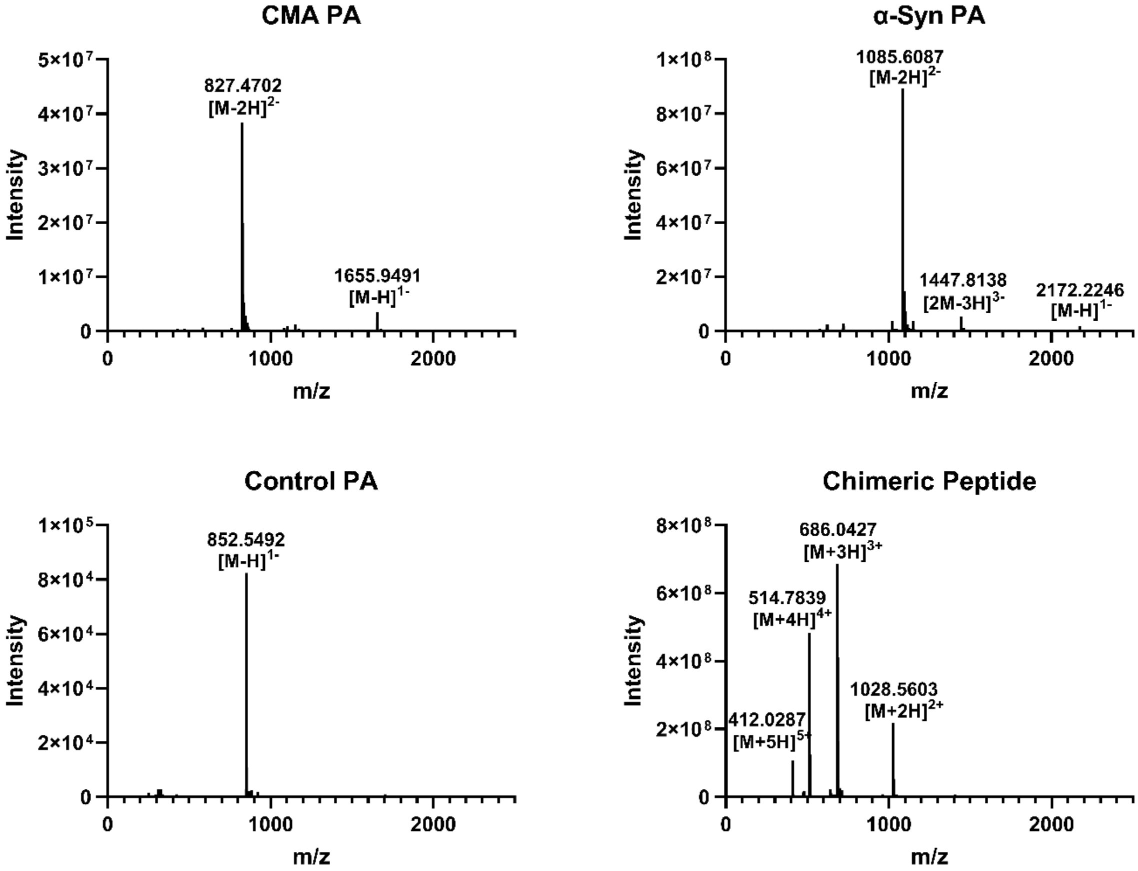

**Figure S1. Electrospray ionization mass spectrometry analysis of purified peptides.** All samples were prepared at 10 µg/mL in 50:50 $H_2O$:ACN containing either 0.1% $NH_4OH$ (PAs) or 0.1% Formic Acid (Chimeric Peptide).

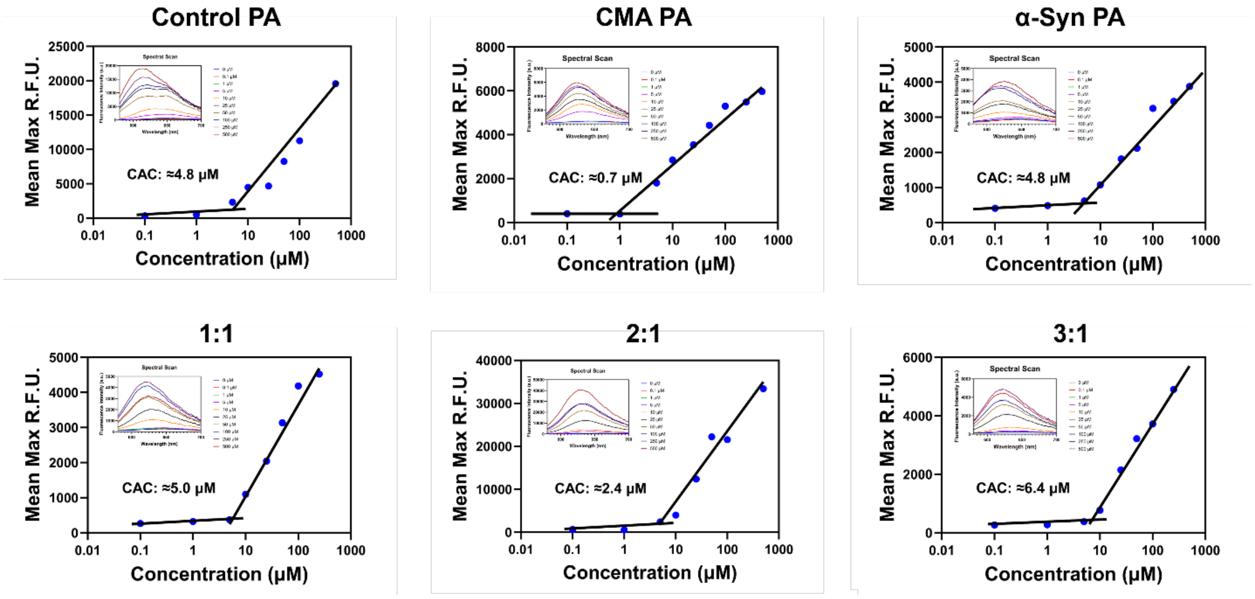

**Figure S2. Critical aggregate concentration determination by Nile red assay**.

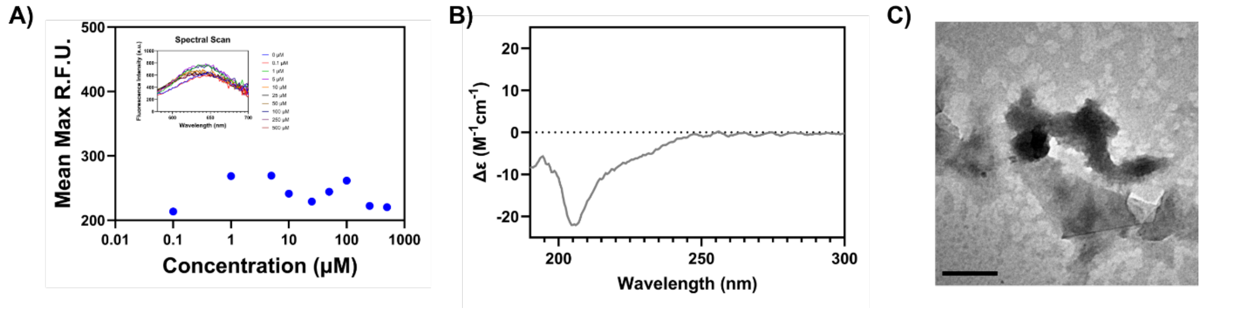

**Figure S3. Nanostructure Characterization of Chimeric Peptide.** A) A Nile red assay indicated no critical aggregation concentration in aqueous solutions consisting of concentrations of up to 1000 μM of chimeric peptide. B) Contrary to PA assemblies, CD did not indicate the presence of beta-sheet formation. Instead, the chimeric peptide exhibited a spectra more closely resembling a characteristic random coil curve. C) TEM Imaging illustrated a lack of ordered, elongated nanofiber formation by the chimeric peptide. Scale bar is 250 nm.

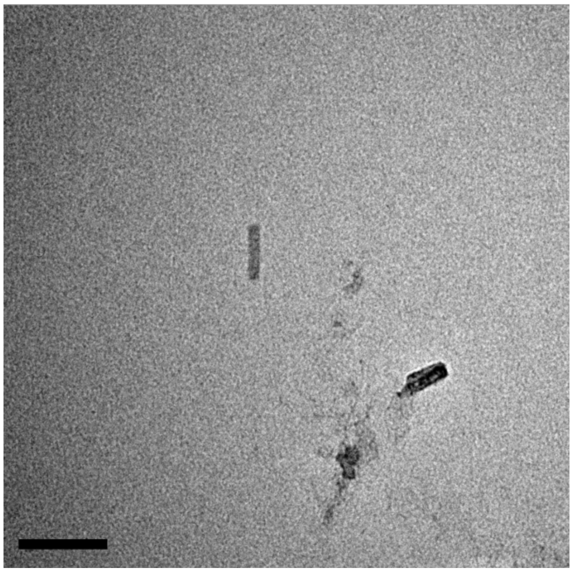

**Figure S4. TEM Imaging of 1:3 (CMA Pa, α-Syn PA, mol%) co-assembly.** TEM Imaging illustrating a lack of ordered, elongated nanofiber formation a 1:3 co-assembly (CMA PA: α-Syn PA, mol%). Scale bar is 100 nm.

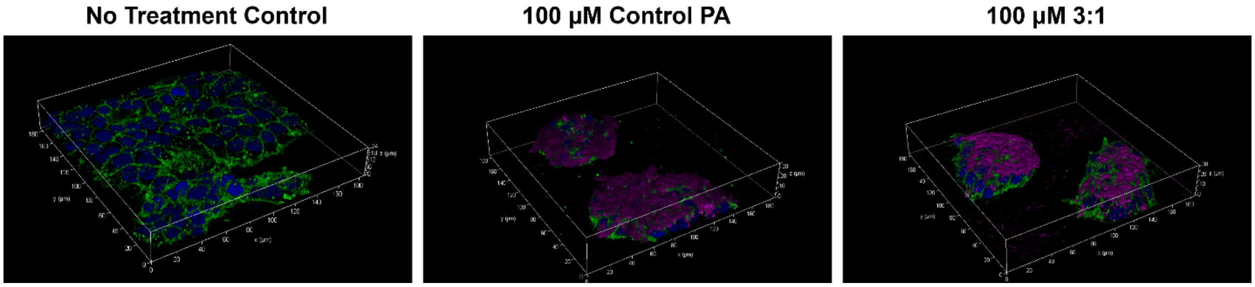

**Figure S5. Confocal microscopy of PA cellular internalization Z-stacks.**

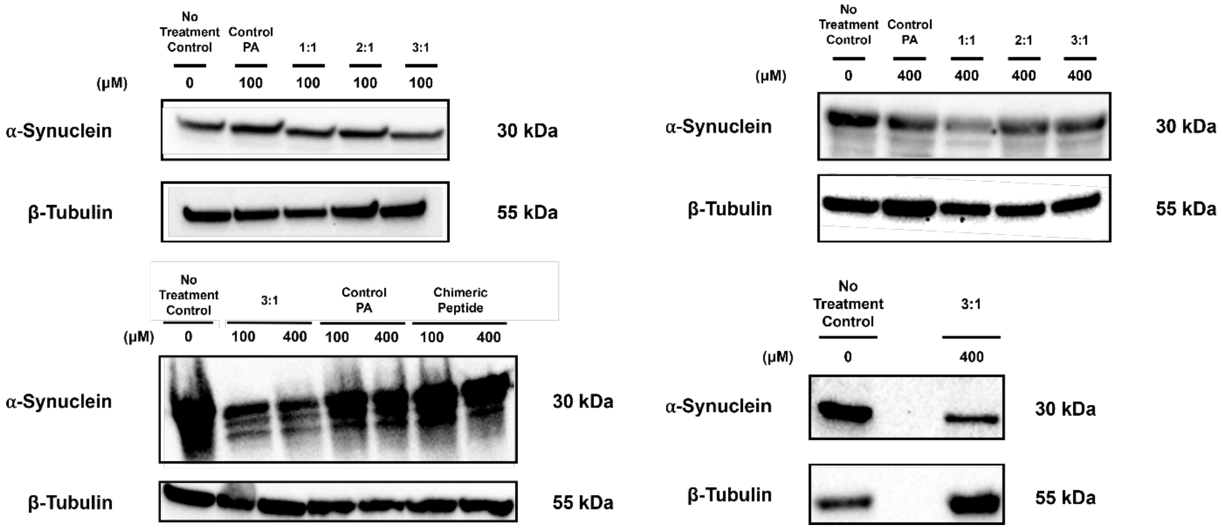

**Figure S6. Additional western blot experiments performed followed TPD treatments.**
